# Meta-azotomics of engineered wastewater treatment processes reveals differential contributions of established and novel models of N-cycling

**DOI:** 10.1101/2020.08.25.229054

**Authors:** Mee-Rye Park, Medini K. Annavajhala, Kartik Chandran

**Author notes:** Joint BioEnergy Institute, Emeryville, CA 94608, USA. Biological Systems and Engineering Division, Lawrence Berkeley National Laboratory, Berkeley, CA 94720, USA. Department of Medicine, Columbia University Medical Center, New York, NY 10032, USA.

## Abstract

The application of metagenomics and metatranscriptomics to field-scale engineered biological nitrogen removal (BNR) processes revealed a complex N-cycle network (the meta-azotome) therein in terms of microbial structure, *potential* and *extant* function. Autotrophic nitrification bore the imprint of well-documented *Nitrosomonas* and *Nitrospira* in most systems. However, in select BNR processes, complete ammonia oxidizing bacteria, comammox *Nitrospira*, unexpectedly contributed more substantially to ammonia oxidation than canonical ammonia oxidizing bacteria, based on metatranscriptomic profiling. Methylotrophic denitrification was distinctly active in methanol-fed reactors but not in glycerol-fed reactors. Interestingly, glycerol metabolism and N-reduction transcript signatures were uncoupled, possibly suggesting the role of other carbon sources in denitrification emanating from glycerol itself or from upstream process reactors. In sum, the meta-azotome of engineered BNR processes revealed both traditional and novel mechanisms of N-cycling. Similar interrogation approaches could potentially inform better design and optimization of wastewater treatment and engineered bioprocesses in general.

## Introduction

The microbial ecology of engineered biological wastewater treatment plants (WWTPs) is key to the transformation of pollutants and nutrients therein and is in turn shaped by the type of wastewater and the WWTP operating conditions. An understanding of microbial communities could thus be crucial to developing and applying sustainable and efficient wastewater treatment technologies (*1*). From this perspective, it is equally important to understand the distribution, abundance and metabolic functions of dominant microbial communities. This is especially true in the case of increasingly complex engineered nutrient removal systems such as biological nitrogen removal (BNR) systems.

With recent advances in high-throughput sequencing technologies (*2*–*6*), our understanding of the phylogenetic and taxonomic classification of activated sludge microbial communities has continued to expand. Nonetheless, the ecology of whole microbial communities is thus far largely derived from studies on single gene biomarkers (at the DNA level). Little information is available regarding the relationships between the community of active microbes assessed and their activity in engineered BNR systems, which can be obtained by targeted sequencing of mRNA. Moreover, we lack an integrated understanding regarding the possibility and extent to which community gene expression in engineering BNR systems is influenced by process operating conditions, such as alternating oxic-anoxic transitions, which are integral to nitrogen removal through nitrification and denitrification.

Accordingly, in this study, we explored the potential and extant function of several field-scale BNR processes using a combination of metagenomics (whole-genome, targeting the DNA pool) and metatranscriptomics (targeting the messenger RNA, mRNA pool) approaches, respectively. The specific objectives were to: (1) understand metagenomics-derived whole community structure and potential function from full-scale BNR plants representing a broad spectrum of configurations and operating conditions, (2) characterize the metatranscriptomic responses pertaining to nitrogen and external carbon metabolism in response to anoxic-aerobic transitions within a given activated sludge process, and (3) explore and elucidate extant models and pathways involved in overall community N- and related organic carbon-cycling.

## Results and discussion

### Unsupervised cluster analysis of community structure and function obtained from shotgun metagenomics

Prior to further detailed analysis or interpretation, the metagenomes from the BNR processes were first subjected to unsupervised clustering to derive broad inferences on any relationships amongst them, in terms of microbial structure and functional potential (Fig. S1). Based on hierarchical cluster analysis at the genus level, three main clusters were apparent. Cluster 1 was comprised of the two glycerol-fed processes, AT-3 and AT-13, which clustered together with high similarity of 70.7%, along with the aeration tank at York River, YRARE (operated without any external carbon) with a lower similarity (52.5%). Of the processes sampled, the YRARE process uniquely operated without an additional carbon source. The second cluster (with a 55.5% similarity) was comprised of DCW, NP, and YRDENI processes, which are all methanol-fed. However, the YRDENI was different from the DCW-NP group, presumably associated with its operation as a denitrification filter, not significantly influenced by the primary effluent composition or suspended growth activated sludge. Finally, the SHARON process was substantially different from the remaining processes, likely associated with its distinct isolated model of treating sidestream centrate at a controlled higher temperature of approximately 33°C. Therefore, although cluster analysis does not provide any mechanistic knowledge into the structure-function-operations links amongst the processes sampled, some qualitative insights and patterns were already apparent.

### Metagenomics-derived overall community structure

Although each process operated under different reactor configurations and operational conditions (e.g., temperature, external carbon source, nitrogen and carbon loads), five genera were abundantly observed (>0.5%) in all seven reactors. These included *Acidovorax, Pseudomonas*, and *Burkholderia*; ammonia-oxidizing bacteria (AOB) such as *Nitrosomonas* spp.; and polyphosphate-accumulating organisms (PAO)/denitrifiers such as *Ca*. Accumulibacter (Fig. S2). The other frequently observed and abundant genera in six samples included *Thauera*, *Azoarcus*, *Bradyrhizobium*, *Dechloromonas*, *Flavobacterium*, *Flavihumilbacter*, *Terrimonas* as well as methanotrophic bacteria such as *Methylibium*; and iron-oxidizing bacteria (IOB) such as *Sediminibacterium* (Fig. S2). Most of these genera are reportedly widespread in activated sludge communities (*7*).

### Bacterial and archaeal community composition involved in autotrophic nitrification

*Nitrosomonas* spp. were the most abundant AOB (0.8-4.3% of the microbial communities sampled) detected in this study (Fig. S2). Among the AOB, *Nitrosomonas* are metabolically versatile and have been detected in a wide range of natural and engineered environments subject to different extant ammonia concentrations and operating conditions (*8*, *9*). After *Nitrosomonas* (63-99% of the total AOB sequences), *Nitrosospira* (0-32% of the total AOB sequences) and *Nitrosococcus* (0-5% of the total AOB sequences) were the most abundantly detected AOB but not in every process sampled (Fig. 1(A)).

**Fig. 1.**
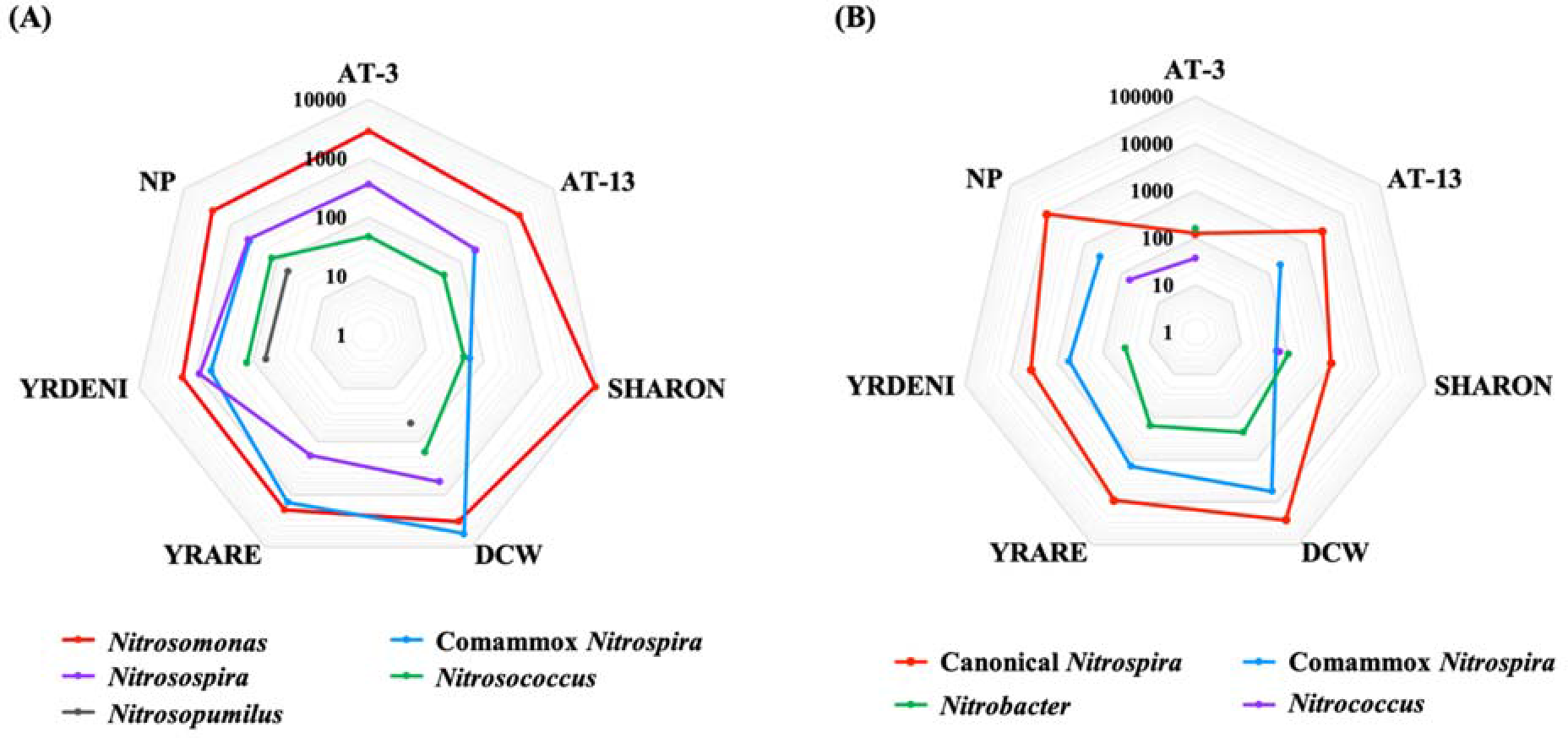
Comparison of (A) AOB, AOA and (B) NOB in each sample. Indicated values are sequence read numbers in logarithmic scale. Comammox *Nitrospira* are shown in both (A) and (B) since comammox *Nitrospira* are functionally represented in both ammonia and nitrite oxidation.

Among NOB, *Nitrospira* (including both canonical and comammox *Nitrospira*) were abundantly detected (1.3-13.5% of the microbial communities sampled, Fig. S2) in AT-13, DCW, YRARE, YRDENI and NP (all mainstream BNR processes), and to a lesser extent (0.1-0.3% of the microbial communities sampled, Fig. S2) in AT-3 and SHARON (both sidestream BNR processes). *Nitrospira*-related NOB have higher affinity coefficients for nitrite and oxygen affinities (*10*, *11*). Moreover, a previous study estimated that the growth stoichiometry and thermodynamic efficiency for *Nitrospira* were higher than for *Nitrobacter* (*11*). The higher electron capture efficiency and biomass yields of canonical *Nitrospira* likely contribute to their increased competitiveness and widespread presence in mainstream engineered BNR systems with limiting extant substrate (including nitrite and oxygen) concentrations rather than sidestream BNR systems.

Specifically, in the mainstream DCW process, comammox *Nitrospira* (15.2% of the total sequences of AOB and NOB) were even more abundant than *Nitrosomonas* related AOB (9.0% of the total sequences of AOB and NOB) (Fig. 1(A)). Overall, the metagenome reads of canonical *Nitrospira* (39-96% of the total NOB sequences) systematically exceeded those of comammox *Nitrospira* (0-17% of total NOB sequences) and *Nitrococcus* (0-11% of total NOB sequences) in all reactors, and exceeded *Nitrobacter* reads (0-50% of the total NOB sequences) in all reactors except AT-3 (Fig. 1(B)). Whereas canonical *Nitrospira* and *Nitrobacter* have traditionally been regarded as the main NOB in WWTPs (*12*), the results of this study show the near preponderance of comammox *Nitrospira* therein, albeit at lower levels than canonical *Nitrospira* spp. (Fig. 1(B)). Recently, the ubiquity of comammox *Nitrospira* in WWTPs across the globe has also be shown through shotgun metagenomics (*2*). The discovery of comammox has dramatically changed our understanding of the microbial nitrogen cycle in engineered and natural environments (*13*, *14*). As such, future characterization and optimization efforts of BNR processes need to include the contributions of comammox to N-cycling therein.

Ammonia oxidizing archaea (AOA) related only to the genus *Nitrosopumilus* were detected in DCW, YRDENI, and NP, although at much lower abundance than AOB (Fig. 1(A)). The contribution of AOA to ammonia oxidation in engineered BNR processes is still not conclusive (*15*, *16*). Our results also suggest a more substantial role for bacterial than archaeal nitrification in the processes sampled.

### Impact of external carbon source on chemoorganoheterotrophic denitrifying community structure

Expectedly, the methanol-fed processes including DCW, YRDENI and NP particularly selected for methylotrophic denitrifying populations related to *Methylotenera*, *Hyphomicrobium* and *Methylophilus* (Fig. S2). In contrast diverse non-methylotrophs were present in the glycerol-fed reactors such as AT-3, AT-13 and SHARON (Fig. S2). Previously, based on ^13^C-DNA SIP, *Hyphomicrobium* spp. and *Methyloversatilis* spp. were identified as the predominant active fractions in methylotrophic denitrification in activated sludge (*17*). Another study reported that ^13^C glycerol-assimilating denitrifiers were related to *Comamonas, Bradyrhizobium* and *Tessaracoccus* (*18*). Although some subgroups of methylotrophic denitrifiers can grow on a wide range of carbon compounds (*19*, *20*), methylotrophs were largely restricted to methanol-fed processes in this study, strongly underscoring the link between process operations and the supported microbial community structure and function.

### Singular contribution of internally produced sidestreams to archaeal diversity in corresponding treatment processes

Interestingly, *Methanolinea* and *Methanosaeta* among archaea were only abundantly represented in AT-3 and SHARON, both of which treat anaerobic digestion centrate (Fig. S2). The higher prevalence of archaeal sequence reads in the separate centrate treatment processes aligns with archaeal methanogenesis in the upstream digesters, from which the centrate fed to these two processes is derived. *Methanolinea* and *Methanosaeta* represent hydrogenotrophic and acetoclastic methanogens that convert hydrogen gas and acetate to methane gas, respectively. This finding underlines the role that singular influent streams (*in casu*, anaerobic digester supernatant or centrate) have in shaping the community structure (and potentially function) of downstream process reactors treating such streams.

### Meta-transcriptomics-derived extant community function: Nitrogen and organic carbon metabolism in anoxic relative to aerobic conditions

Anoxic-oxic cycling is integral to the design and operation of typical BNR process reactors (*21*). Broadly, aerated zones are designed to support aerobic oxidation of organic carbon and nitrification and non-aerated mechanically mixed zones are designed to support denitrification with or without the addition of external electron donors, including organic compounds. Anaerobic zones can further be integrated into BNR processes to effect removal of phosphorus as well (*21*). In this study, we evaluated spatial profiles of mRNA concentrations of genes coding for nitrification, denitrification and external carbon metabolism (glycerol and methanol) pathways to further understand *in-situ* microbial activity under anoxic conditions versus oxic conditions specifically for the SHARON, NP and DCW processes (Fig. 2). This approach draws from previously observed parallels between the expression of select genes and whole-cell activity measures (*22*–*27*). For nitrification, we spatially profiled the genes coding for ammonia oxidation to hydroxylamine (ammonia monooxygenase, *amo*), hydroxylamine oxidation to nitrite (hydroxylamine oxidoreductase, *hao*), and nitrite oxidoreductase (*nxr*). For chemoorganoheterotrophic denitrification we examined the spatial expression profiles of *nar*/*nap*, *nir*, *nor* and *nos*, which are associated with nitrate reductase, nitrite reductase, nitric oxide reductase and nitrous oxide reductase, respectively (*28*). We expected that the BNR community would reduce the expression of nitrification genes and increase the expression of denitrification genes in response to alternating anoxic conditions to aerobic conditions.

**Fig. 2.**
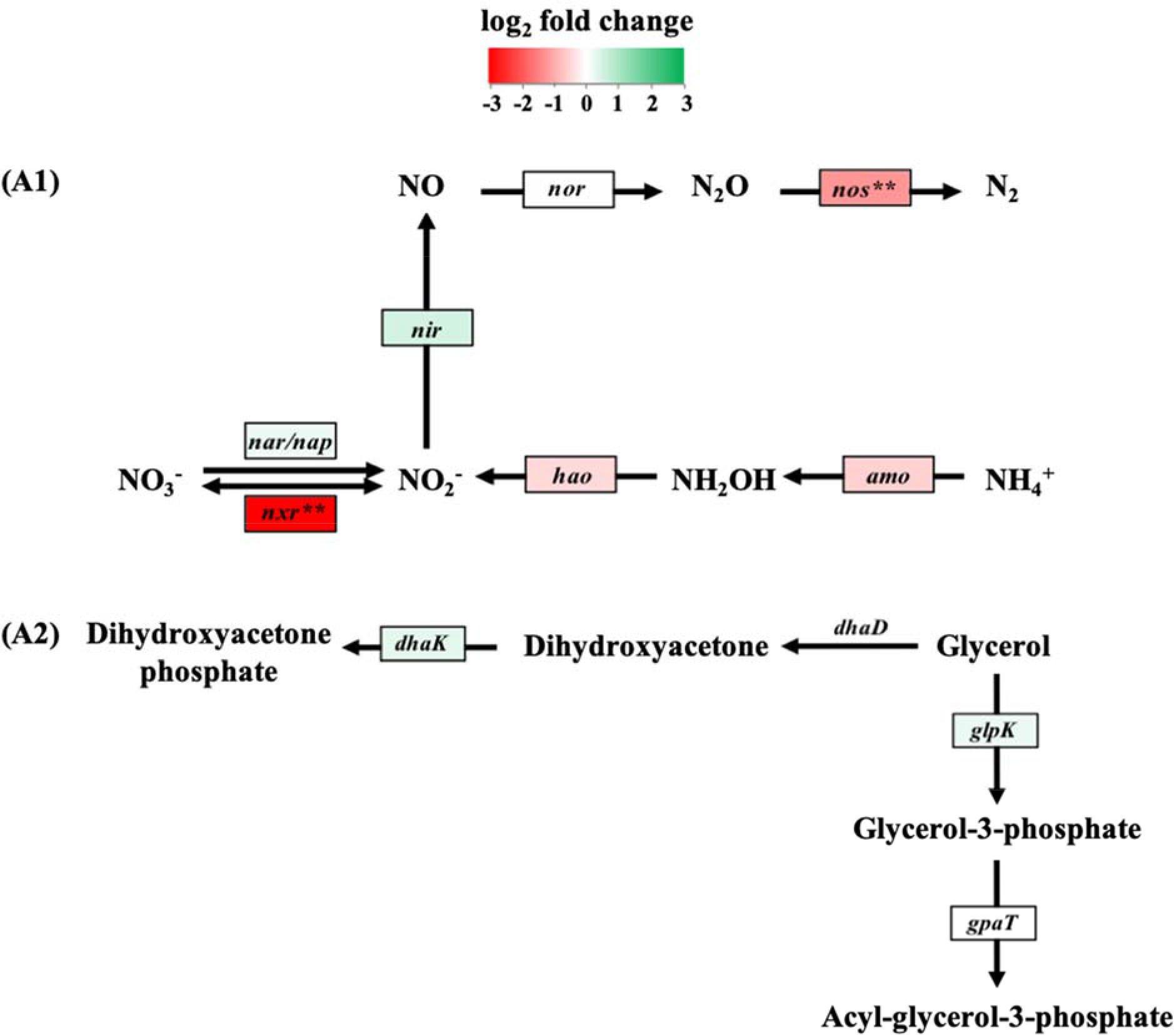
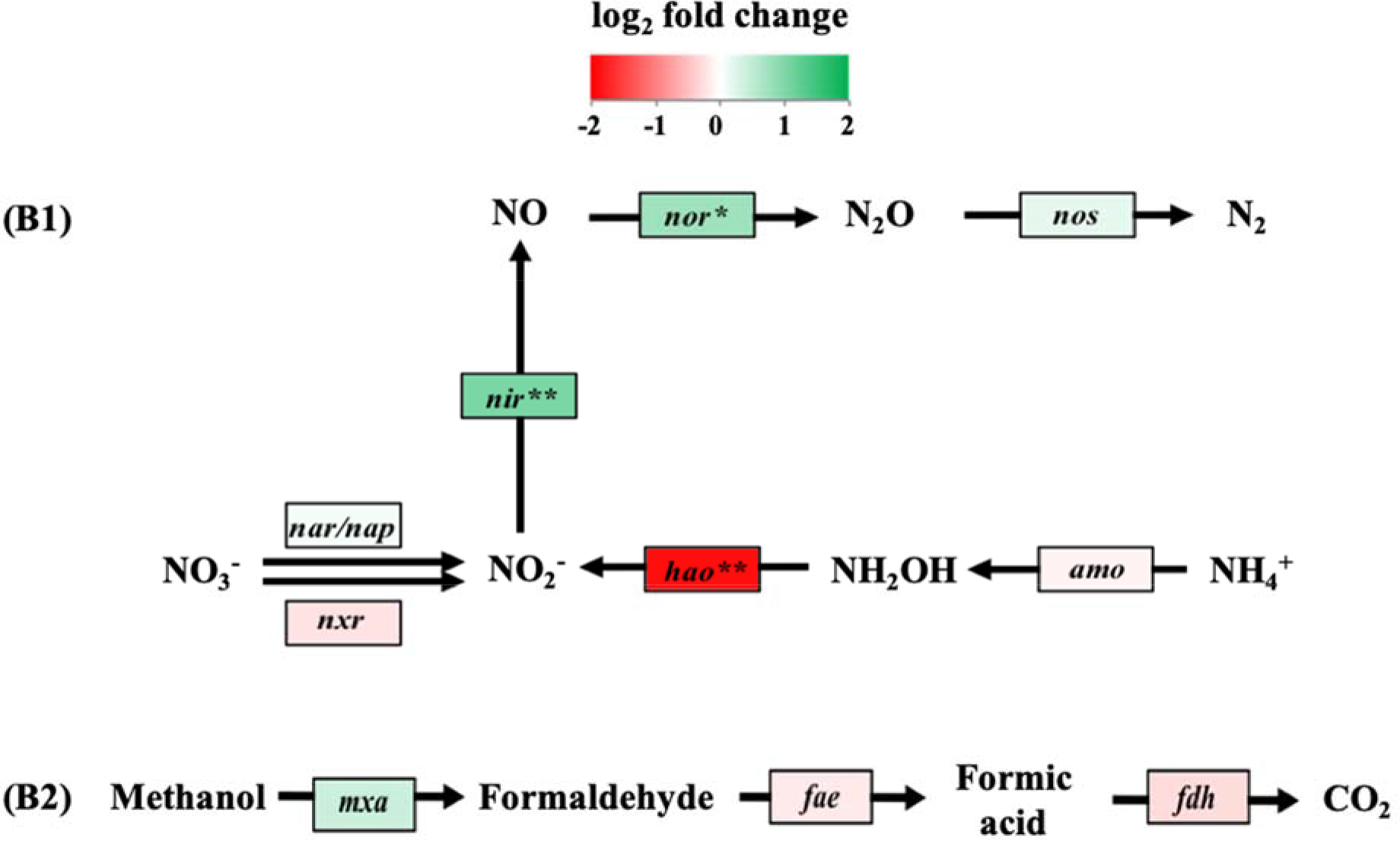
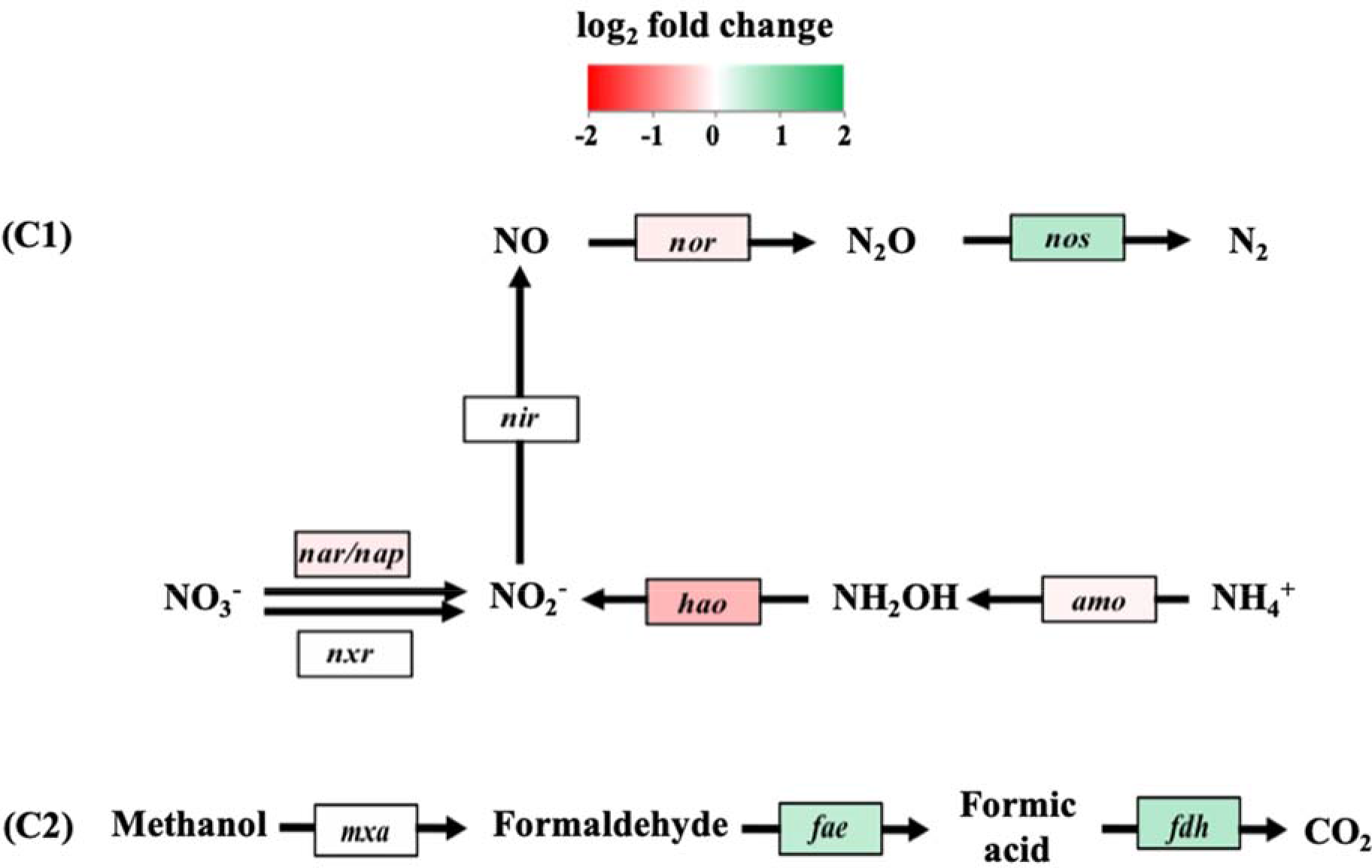
Differential expression of genes involved in nitrogen and organic carbon cycling. (**A1**) nitrification, denitrification and (**A2**) glycerol oxidation pathways encoded by SHARON metatranscriptome; (**B1**) nitrification, denitrification and (**B2**) methanol oxidation pathways encoded by NP metatranscriptome; (**C1**) nitrification, denitrification and (**C2**) methanol oxidation pathways encoded by DCW metatranscriptome. Color intensity represents fold change based on the log-fold change in mRNA transcript levels of anoxic condition relative to aerobic condition. Gene expression was relativized by RPKM values. Symbols reflect statistically signinficant changes in gene expression: \*\**p* < 0.05 and \**p* < 0.10.

In the SHARON process, as expected, the transcript levels of nitrification genes (*amo, hao*, and *nxr*, respectively) reduced as the biomass transitioned from the aerobic to anoxic zones relative to aerobic conditions (denoted in red color, Fig. 2(A1) and Table S2). In contrast, but again in line with expectation, for denitrification with glycerol as carbon source, the expression of most nitrogen reductase genes were higher under anoxic conditions than under aerobic conditions (denoted in green color, Fig. 2(A1) and Table S2). Unexpectedly, *nos* transcription levels showed a decrease under anoxic conditions. In general, NOS is more sensitive to a variety of environmental factors than the other upstream reductases (*29*, *30*). However, the specific factors contributing to its decreased expression under anoxic conditions cannot be conclusively obtained from the overall system-approach followed herein. More detailed and high-resolution (chemical and molecular) measurements will likely be more helpful in this regard. With respect to organic carbon-cycling, the first step of glycerol oxidation involves glycerol kinase (coded by *glpK*) or glycerol dehydrogenase (coded by *dhaD*) and the expression of both genes has been explored as a biomarker of glycerol-based denitrification activity (*31*, *32*). Here, the transcript levels of *glpK* gene displayed increased under anoxic relative to oxic conditions, whereas *dhaD* transcripts were not detectable (Fig. 2(A2) and Table S2). These results possibly suggest that the denitrifiers in the SHARON process preferentially employ the glycerol kinase (*glp*) pathway instead of the *dhaD* encoded pathway. Added insights into the uncoupling of glycerol metabolism and N-reduction activity were also revealed using the functional gene transcripts and these are described below for the SHARON process, *linking structure and expressed function*.

In the NP process, the transcript levels of *amo, hao*, and *nxr* showed expected decreases under anoxic conditions relative to aerobic conditions (Fig. 2(B1) and Table S2). Similarly, for denitrification, the expression of genes coding for nitrogen reductase pathways as well as methanol oxidation were higher in the anoxic zones than in the aerobic zones (Fig. 2(B1)). However, the expression of formaldehyde activating enzyme (*fae*) and formate dehydrogenase (*fdh*), which catalyses the metabolism of formaldehyde and formic acid respectively, were unexpectedly lower in the anoxic zones (Fig. 2(B2) and Table S2). Even such unexpected trends are quite revealing suggesting reaction-specific control within the denitrification cascade (of both carbon and nitrogen pathways) under the conditions imposed in engineered BNR processes.

In the DCW process, which entails nitrification followed by methanol-fed nitrification, the transcript levels of the nitrification coding genes, *amo*, *hao*, and *nxr* were lower in the anoxic zone that followed the aerobic zone, as expected (Fig. 2(C1) and Table S2). For denitrification, the increased expression of methanol oxidation genes in the anoxic relative to oxic zones was also in line with expectation (Fig. 2(C2) and Table S2). However, the expression of *nos* showed an unexpected increase in the anoxic zone (Fig. 2(C1)).

Notwithstanding some specific deviations (*nos* expression in the SHARON and DCW processes), it is remarkable to observe that even in highly variable full-scale WWTPs, there was good overall correspondence between i*n-situ* transcript profiles of denitrification (both nitrogen and carbon transformation pathways) and process operations. Exploring this correspondence further could have the added utility of using *in-situ* transcript profiles (spatial or temporal) as useful diagnostics for process health.

### Transcription profiles corresponding to nitrification, denitrification and external carbon metabolism – linking microbial structure and expressed function

From a microbial “functional” perspective, the relative contributions of the different microbial communities within the SHARON, DCW and NP processes were further analyzed by targeting key functional gene transcripts (based on RPKM) of nitrification, denitrification and organic carbon oxidation pathways. Overall, the transcript pools for both nitrogen redox transformations and carbon utilization displayed significantly different diversities (Fig. 3). Among nitrifying organisms, *Nitrosomonas* spp. and canonical *Nitrospira* spp. imposed the highest contributions, based on *amo* and *hao* transcripts for ammonia to hydroxylamine to nitrite oxidation and based on *nxr* transcripts for nitrite to nitrate oxidation, in the SHARON and NP processes (Fig. 3(A1) and Fig. 3(B1), respectively). Intriguingly, the contributions of comammox *Nitrospira* to *amo* and *hao* transcript pools were significantly higher (66.3% and 100%, respectively) than for the conventional AOB, *Nitrosomonas* spp. (20% and not detected, respectively, at DCW (Fig. 3(C1)). Canonical *Nitrospira* and comammox *Nitrospira* contributed 79.1% and 15.8%, respectively, to NXR (nitrite to nitrate oxidation) at DCW (Fig. 3(C1)). Based on these transcript and metagenomics profiles, a new model for autotrophic ammonia to nitrate oxidation emerges at DCW, with comammox *Nitrospira* driving ammonia to nitrite oxidation and canonical *Nitrospira* driving nitrite to nitrate oxidation. It has been suggested that comammox *Nitrospira* could be competitive under substrate-limited conditions (*33*). However, based on these results, there appear to be even more complex process-specific drivers for comammox proliferation in field-scale BNR systems, which still merit elucidation. In contrast, the metatranscriptomes in the SHARON and NP processes reflect the more conventional role of *Nitrosomonas* spp. and *Nitrospira* spp. in autotrophic ammonia and nitrite oxidation, respectively.

**Fig. 3.**
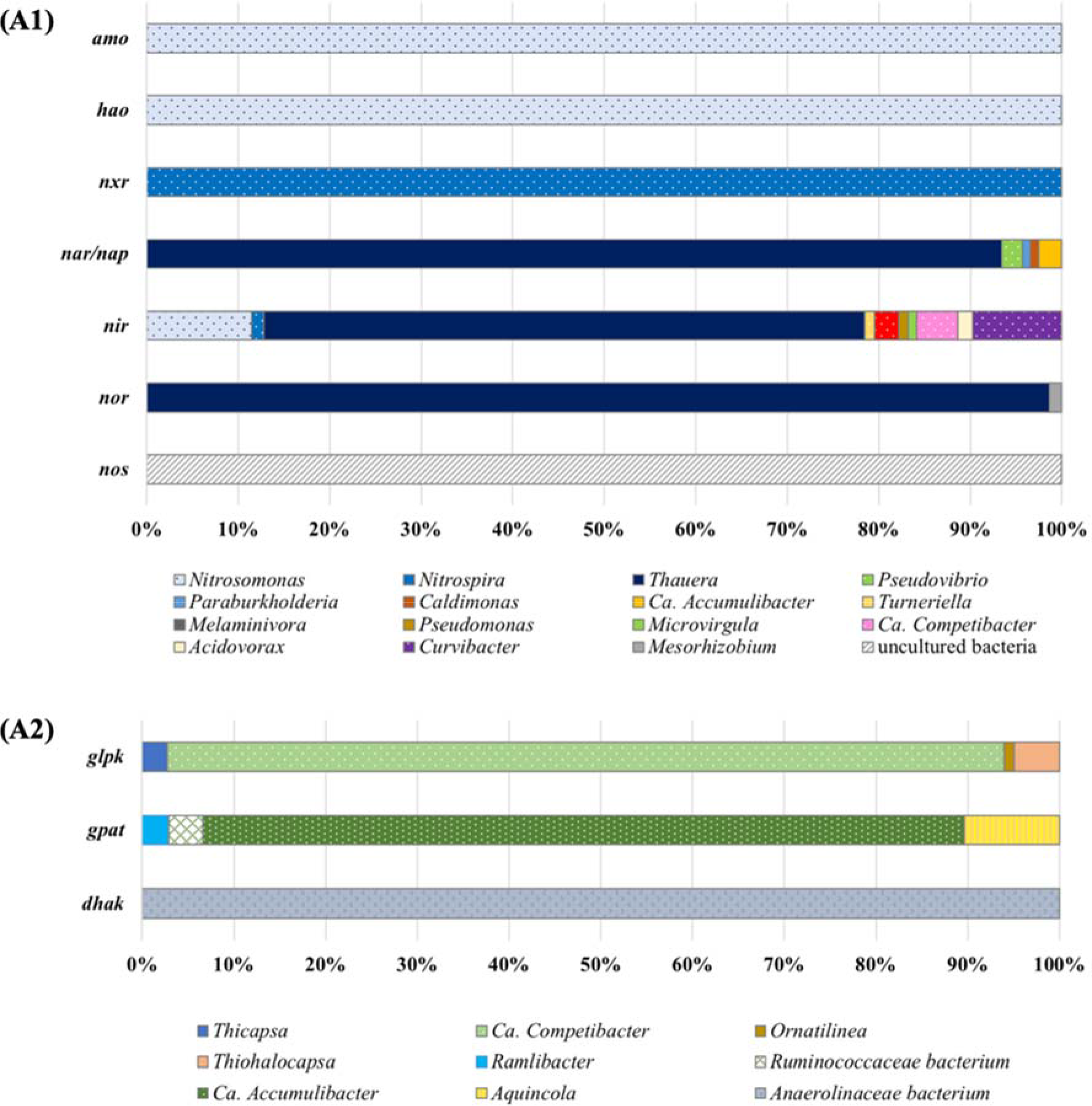
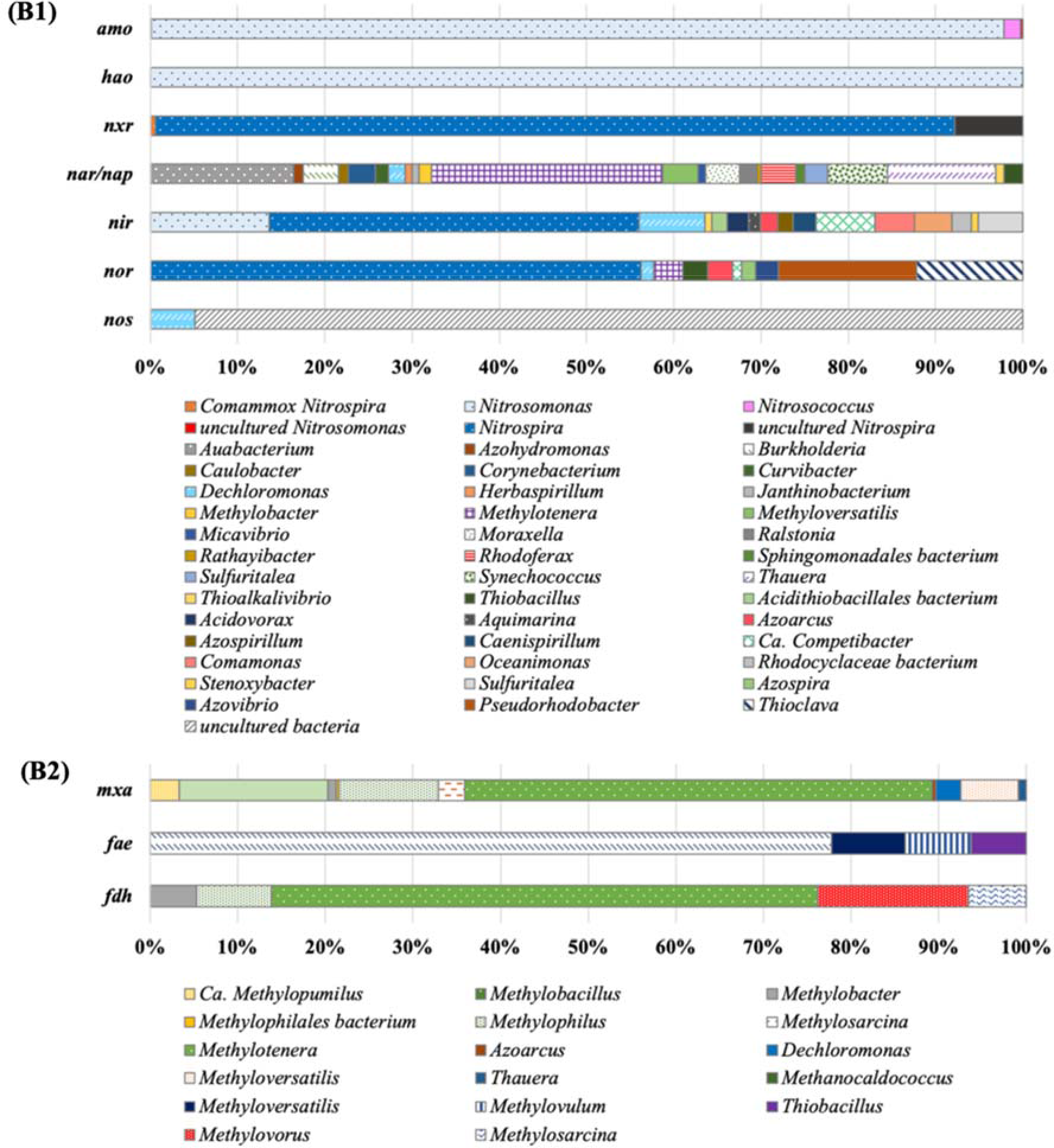
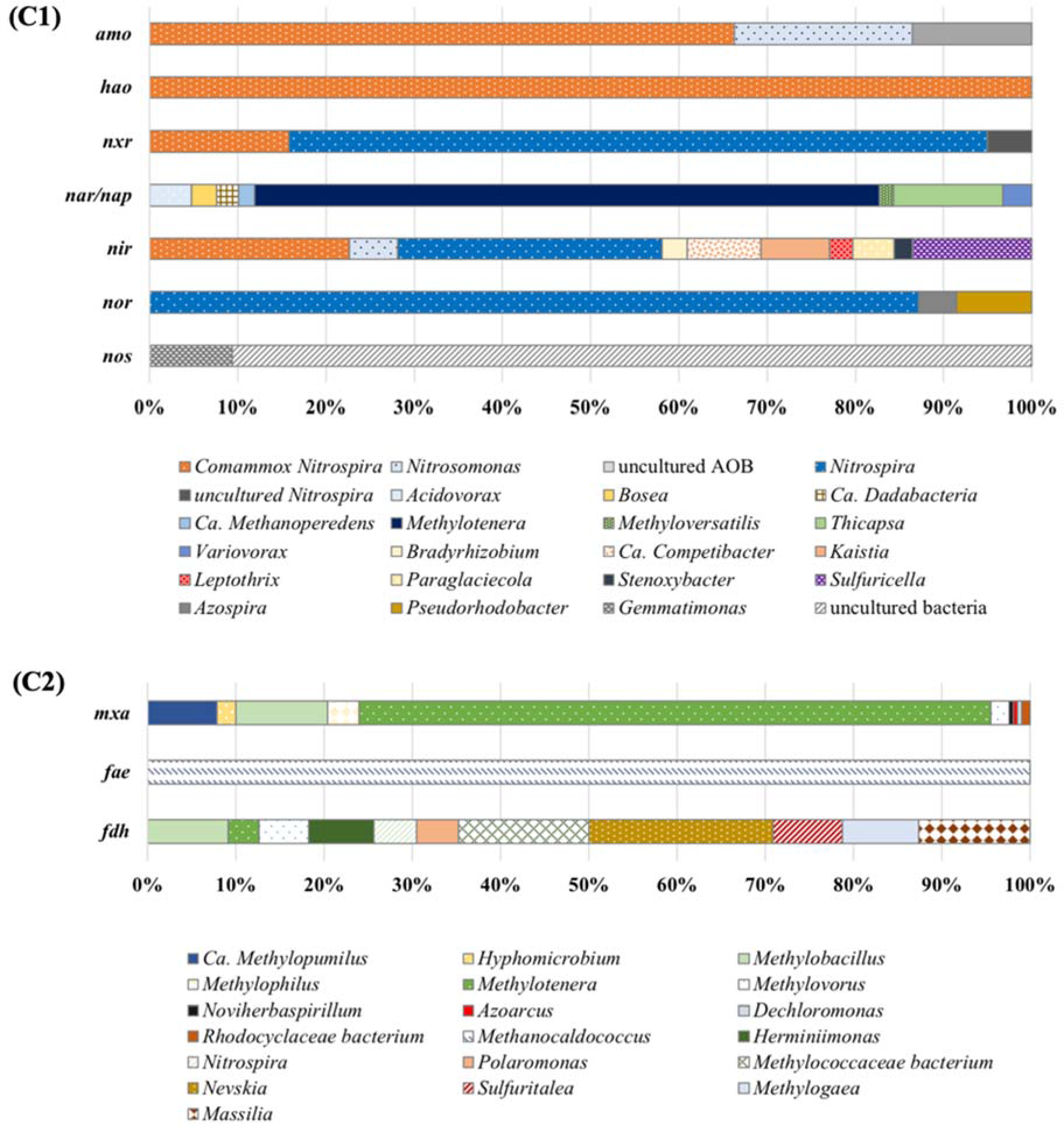
Putative active microorganisms of transcripts encoding enzymes involved in nitrogen and organic carbon cycling. (**A1**) nitrification, denitrification and (**A2**) glycerol oxidation in SHARON metatranscriptome; (**B1**) nitrification, denitrification and (**B2**) methanol oxidation in NP metatranscriptome; (**C1**) nitrification, denitrification and (**C2**) methanol oxidation in DCW metatranscriptome. The relative percentage (%) was calculated from the contributions of different genera in terms of RPKM to total RPKM of each enzyme.

For denitrification, metagenomic reads for methylotrophic denitrifying bacteria were distinctly dominant in methanol-fed reactors, whereas non-methylotrophic denitrifiers were shown in glycerol-fed reactors based on metagenomes. Surprisingly, glycerol metabolism and N-reduction activity were uncoupled herein in terms of functional gene transcripts involved in denitrification and glycerol metabolism. For example, in SHARON, the highest contribution of *nar*, *nir* and *nor* transcripts were from *Thauera*, but the transcription of *glp* was not observed in *Thauera* (Fig. 3(A)). This suggests that *Thauera* likely did not employ glycerol as an external carbon source for denitrification but rather utilized glycerol metabolites, such as glyceraldehyde, produced by other members of the community. For instance, the *glp* mRNA transcripts were observed in other bacteria including *Thiocapsa*, *Ca*. Competibacter, *Ornatilinea* and *Thiohalocapsa*, which employed a glycerol kinase pathway encoded by the *glp* system, but *Thiocapsa, Ornatilinea* and *Thiohalocapsa* did not contribute to the transcript pools of the nitrogen reductase genes. Solely *Ca*. Competibacter, which has been implicated in glycogen production and storage, expressed *nir* mRNA transcripts in parallel with *glp* transcripts, indicating coupled glycerol-based nitrite reduction. These observations raise the prospect of an intricate network of interactive carbon and nitrogen exchanges in engineered BNR processes, even with a relatively ‘simple’ organic electron donor such as glycerol. Such exchanges and interactions need to be integrated into mechanistic process modeling and operations.

In the methanol-fed reactors (NP and DCW), the transcripts of *nar* and *mxa* bore the imprint of the documented methylotrophs, *Methylotenera* spp. (Fig. 3(B) and Fig. 3(C)) *Methylotenera* spp. were also the most abundant methylotrophic bacteria in metagenomes of both NP and DCW. Besides *Methylotenera* spp. in NP and DCW, diverse denitrifiers that utilized neither methanol nor methanol intermediates were observed based on mRNA transcripts of denitrification genes, which were likely correlated with denitrification fostered by other carbon sources in the reactors. Interestingly, at NP and DCW, the highest transcript levels of *nir* (coding for NO_2_^-^ reduction to NO, 42% and 29%, respectively) and *nor* (coding for NO reduction to N_2_O, 56% and 87%, respectively) corresponded to *Nitrospira* spp. (Fig. 3(B1) and Fig. 3(C1) This is similar to recent observations of NOB-related nitrogen reductase transcripts in nitrification reactors subjected to repeated anoxic-aerobic cycling (*3*) and introduces the added contribution of some NOB to NO and N_2_O production in engineered BNR processes.

In sum, the meta-azotome of a diverse set of engineered BNR processes reveals a combination of established and novel knowledge pertaining to the structure-function-activity of N- and C-cycling by mixed microbial communities therein. At the structural and ‘potential’ functional level, the metagenomics approach followed herein vastly expands the insights possible through traditional targeted profiling of complex microbial communities. The advantages of such systems level interrogation are especially pronounced for microbial functionality, which cannot be necessarily linked to known genotypes (for instance, chemoorganoheterotrophic denitrification or comammox). Notably, community metatranscriptomics results offer insights into community gene expression profiles as the microbial populations cycle between aerobic and anoxic environments and even highlight the impact of un-controlled or non-ideal operating conditions. Thus, a combination of metagenomics and metatranscriptomics can reveal the structure, function, and activity of diverse microbial communities and enable more informed characterization of engineered microbial processes.

## Materials and methods

### Description of process reactors at wastewater treatment plants and sample collections

The WWTPs interrogated included a broad spectrum of treatment processes and configurations (Table S1). These included the following:

1. The AT-3 process, located at the 26th Wards WWTP in New York, NY, treats anaerobic digester sludge centrate (Fig. S3). The process receives glycerol for enhancing denitrification and is connected to the mainstream treatment processes through a return activated sludge stream (*34*).
2. The AT-13 process at the Wards Island WWTP in New York, NY is a four-pass step-feed BNR process treating mainstream sewage and receives glycerol for enhanced denitrification (Fig. S4).
3. The SHARON process at the Wards Island WWTP, NY treats anaerobic digestion centrate through partial nitrification (conversion of influent ammonia to nitrite) followed by glycerol-based denitrification. The SHARON process differs from the in AT-13 process that is it is not connected to the mainstream BNR processes at Wards Island WWTP (Fig. S5).
4. DC Water’s (DCW) Blue Plains advanced wastewater treatment plant is operated as a two-sludge system consisting of upstream BOD removal system followed by a methanol-fed nitrification-denitrification (NDN) system. Given the focus on engineered BNR processes in this study, the NDN process was the subject of interrogation herein (Fig. S6).
5. The aeration tank (YRARE) at York River treatment plant is configured as a step-feed aeration process, which receives mainstream wastewater and is geared to achieve organic carbon oxidation and nitrification (Fig. S7).
6. At the same WWTP are also present denitrification filters (YRDENI), which are fed the clarified effluent stream from the upstream YRARE process and are designed and operated for methanol-based denitrification (Fig. S8).
7. The Nansemond treatment plant (NP) is operated as a 5-stage Bardenpho process to achieve both nitrogen and phosphorus from the influent mainstream wastewater (Fig. S9).

Field biomass samples were collected from the seven process reactors at the five WWTPs for six consecutive months. (Table S1). Samples were collected from each oxic and anoxic zone where additional carbon source was supplemented (Fig. S3 – Fig. S9). Samples for RNA sequencing were additionally treated with RNAprotect (Qiagen, CA) and immediately frozen in dry ice. Upon arrival at Columbia University, the (1 mL) samples were centrifuged at 10000 RPM for 10 mins and stored at −80 °C for next-generation sequencing and further analysis.

### DNA and RNA extraction

Biomass sample pellets were re-suspended in 1 mL 1 x Tris-EDTA buffer solution (Fisher Scientific, MA) and DNA extraction was performed on a consistent total biomass quantity for all samples using DNeasy mini kit with QIAcube (Qiagen, CA). Extracted DNA and RNA in each sample were quantified using dsDNA HS assay kit and RNA HS assay kit on the Qubit 2.0 Fluorimeter (Life Technologies, NY), respectively.

Samples were interrogated at different levels of resolution (Fig. S10). A total of seven samples were used for DNA sequencing, derived from each sample pooled together in equal masses from all aerobic and anoxic zones and six sampling time points in each of the seven reactors. For metatranscriptomic sequencing, total RNA (tRNA) samples from the oxic and anoxic zones in each of the SHARON, NP and DCW reactors gathered over six months were extracted separately using RNeasy mini kits (Qiagen, CA). The quality and quantity of tRNA were assessed using a NanoDrop Lite Spectrophotometer (Thermofisher, MA). The extracts obtained at different sampling times were subsequently combined separately for the oxic and anoxic zones of the three processes resulting in six combined samples. Each of the six samples was processed in duplicate. Consequently, metatranscriptomics results display *in situ* community responses of anoxic-aerobic transitions within a given activated sludge process. Metagenomic and metatranscriptomic library preparation and sequencing were conducted on an Ion Torrent PGM platform.

### Metagenomic library preparation, sequencing and bioinformatics

Genomic DNA libraries were prepared using the NEBnext Fast DNA Fragmentation and Library Prep Set for Ion Torrent (New England Biolabs, MA). Fragments of 450 bp size were separated and extracted with an E-Gel. SizeSelect™ 2% gel (Life Technologies, NY), which were then amplified, purified, and quantified according to the manufacturer’s instructions for the Ion Xpress™ Plus Fragment Library Kit. An Ion Torrent PGM was used with the Hi-Q 400 template preparation and sequencing kit and a 318v2 sequencing chip (Thermo Fisher, MA). All the quality control steps were performed using Bioanalyzer 2100 (Agilent, Santa Clara, CA).

Reads were filtered with minimum average Phred quality score of 20, minimum length of 100 bp, maximum homopolymeric region length of 8 bp using mothur (*35*) to ensure high quality for subsequent data analysis. Filtered reads were aligned using the Diamond (*36*) blastx program with minimum percent identity of 80%, bit score of 20 and maximum e-value of 1e^−10^ against an in-house database of proteins which expanded the NCBI non-redundant (*nr*) protein database with newly available complete ammonia oxidizers (comammox) and anaerobic ammonium oxidizers (anammox) sequences (*2*). The protein alignments were used to calculate reads per million (RPM) for taxonomic assignment. All raw sequence data have deposited in the NCBI BioProject under accession number PRJNA646165.

### Metatranscriptomic library preparation, sequencing and bioinformatics

In order to provide a set of external RNA controls that enable comparison of transcript concentrations from the different processes sampled, external RNA Controls Consortium (ERCC) spike-in control mixes were added to the total RNA from each selected zone according to the manufacturer’s instructions (Ambion, Life Technologies, Product No. 4455352). Subsequently, ribosomal RNA (rRNA) was removed using the Ribo-Zero Magnetic Kit for bacteria (Epicentre, IL). Metatranscriptome library preparation was performed using the Ion Total RNA-Seq Kit v2 (Life Technologies, NY) and samples were barcoded with IonXpress RNA-Seq barcodes (Life Technologies, NY). The resulting cDNA library was assessed for yield and purity using an Agilent High Sensitivity DNA kit for Agilent 2100 Bioanalyzer (Agilent Technologies, CA). Template preparation with the cDNA libraries followed by ISP enrichment was performed using the Ion OneTouch2 system with Ion PGM™ Hi-Q™ OT2 kit (Product No. MAN0010902) and enriched ISPs were sequenced with Ion Torrent 318 chips with 850 flows according to manufacturer’s instructions (Ion PGM™ Hi-Q™ Sequencing Kit, Product No. MAN0009816). Template preparation using the DNA library, followed by ISP enrichment, was performed using the Ion OneTouch2 system as per the manufacturer’s instructions (Ion PGM™ Hi-Q™ OT2 kit, Product No. MAN0010902). The enriched ISPs were sequenced with Ion Torrent 318v2 chip and sequenced according to the manufacturer’s instructions (Ion PGM™ Hi-Q™ Sequencing Kit, Product No. MAN0009816).

Raw reads of metatranscriptome sequences were filtered with average Phred score of 20 and minimum sequence length of 25 bp using mothur software to ensure high quality for subsequent data analysis (*35*). Four filtered metatranscriptome sequences of each sample were combined to conduct de novo assembly. *De novo* assembly of the combined metatranscriptome reads was conducted using IDBA-tran (*37*). Prodigal (*38*) was used to predict protein-coding features on the assembled metatranscriptome contigs. Each filtered metatranscriptome reads were aligned to the protein-coding features of from Prodigal using Bowtie2 package (*39*). The coding regions predicted by Prodigal were annotated against NCBI nr non-redundant protein database (*36*) through the Diamond blastp program with a maximum e-value of 1e^−15^ and a minimum percent identity of 85%. Differential gene expression analysis was conducted using the edgeR package in R to obtain the counts of normalized reads per kilobase per million reads (RPKM) for the genes (*40*). All raw sequence data have deposited in the NCBI BioProject under accession number PRJNA646165.

### Hierarchical cluster analysis

Hierarchical clustering was used to visualize the similarity of microbial community structures at genus level in seven different processes using R software with the Cluster package (*41*). Hierarchical methods are commonly used in ecology and biology and useful because they are not limited to a pre-determined number of clusters and can display the similarity of samples across a wide range of scales (*42*). Bray-Curtis distance was used to quantify the differences in microbial populations and potential functions between seven processes (*42*, *43*).

## Author contributions

MP and KC contributed to conception and design of the study. MP and MKA performed data analyses. MP and KC wrote the manuscript. KC supervised the research.

## Data and materials availability

All data needed to evaluate the conclusions in the paper are present in the paper and/or the Supplementary Materials.

## Supplementary Materials

**Fig. S1.**
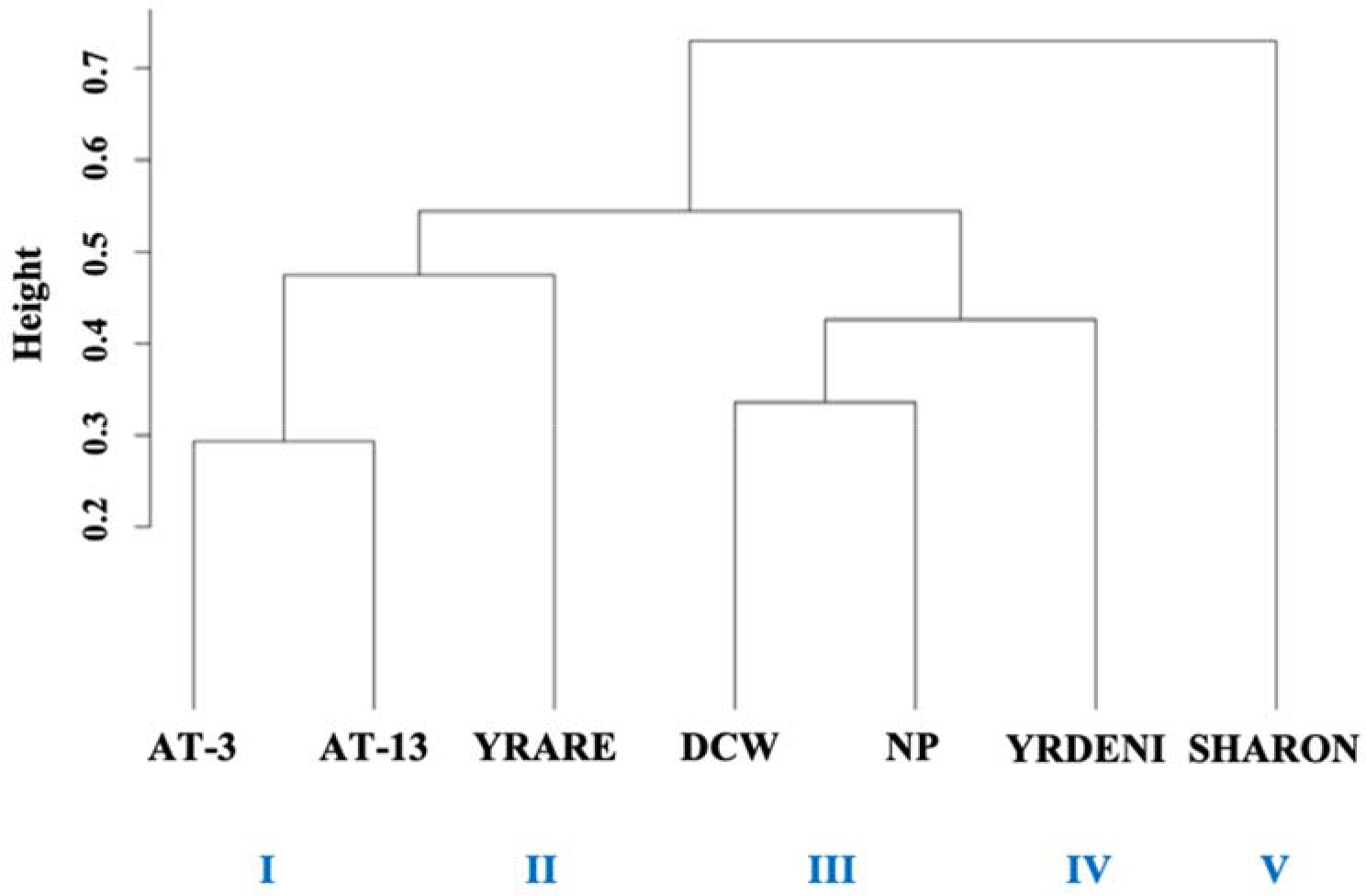
Hierarchical clustering of seven metagenomes at the genus level. Height indicates the level of dissimilarity; branch lengths were computed using Bray-Curtis distance and Ward’s method.

**Fig. S2.**
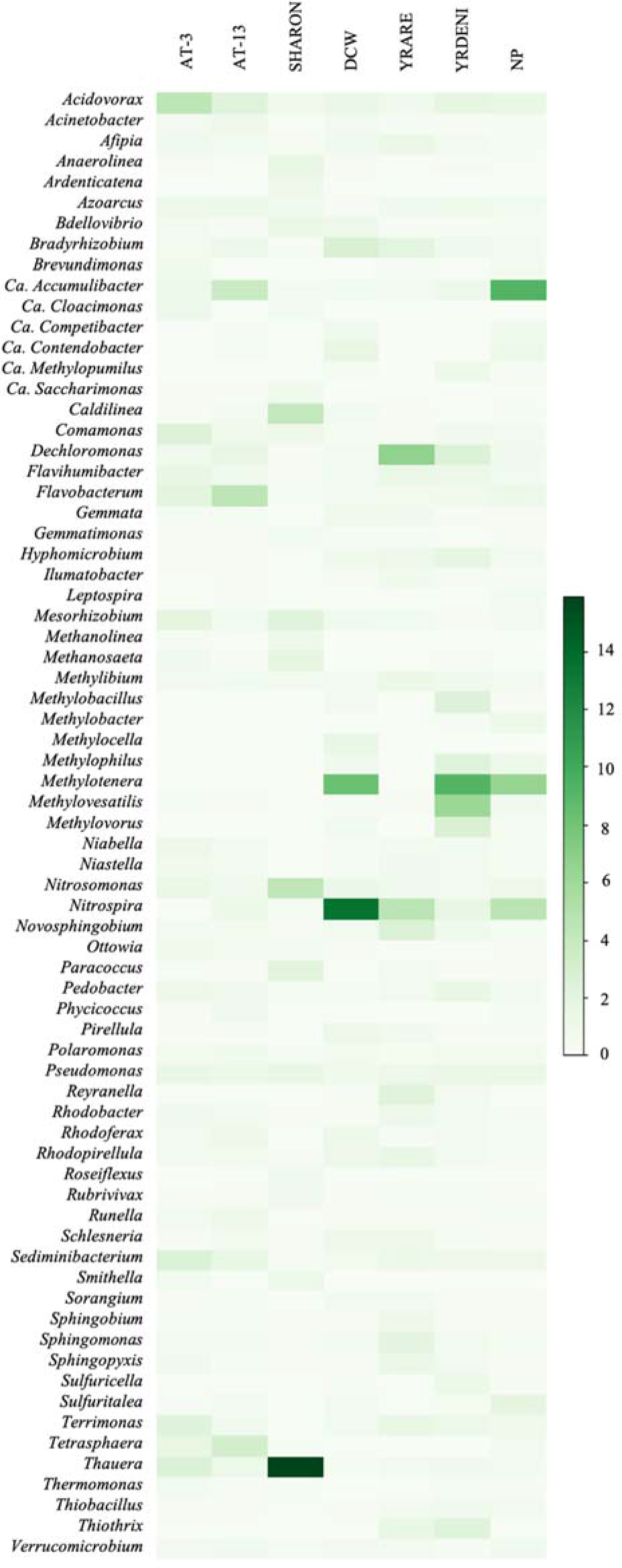
Microbial community composition in samples from seven process reactors. The panel shows relative abundance of the top 20 genera from each sample (a total of 71 genera for all seven samples) as determined from shotgun metagenomic data.

**Fig. S3.**
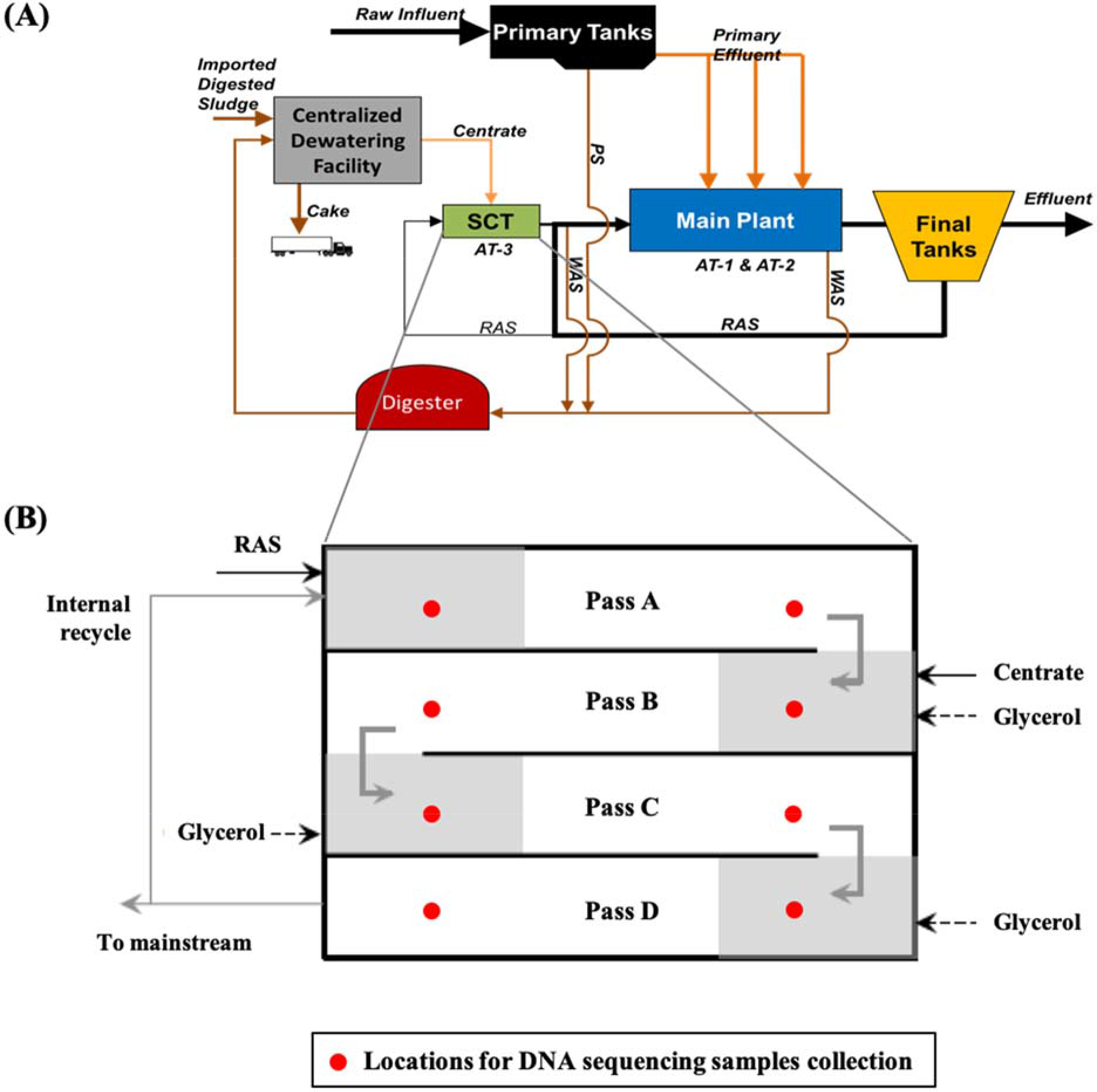
Schematic diagram of (A) 26^th^ Ward WWTP and (B) AT-3 separate centrate treatment (SCT) process with glycerol as the added carbon source. Green arrows indicate the flow of wastewater and shaded areas represent anoxic zones. RAS: return activated sludge. Samples were collected for six consecutive months.

**Fig. S4.**
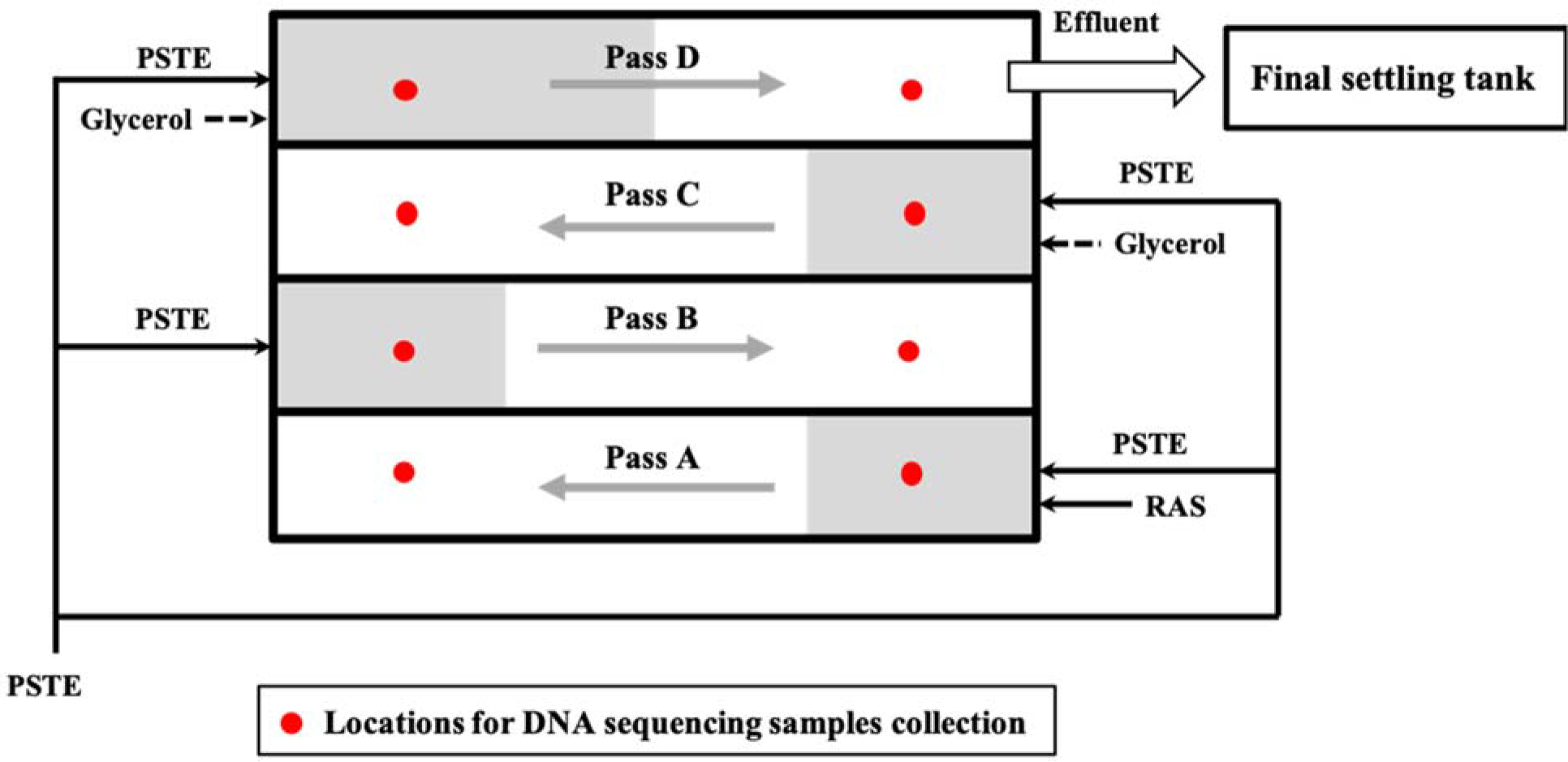
Schematic of AT-13 process at Wards Island WPCP. Shaded areas represent anoxic zones. PSTE: primary settling tank. Samples were collected for six consecutive months.

**Fig. S5.**
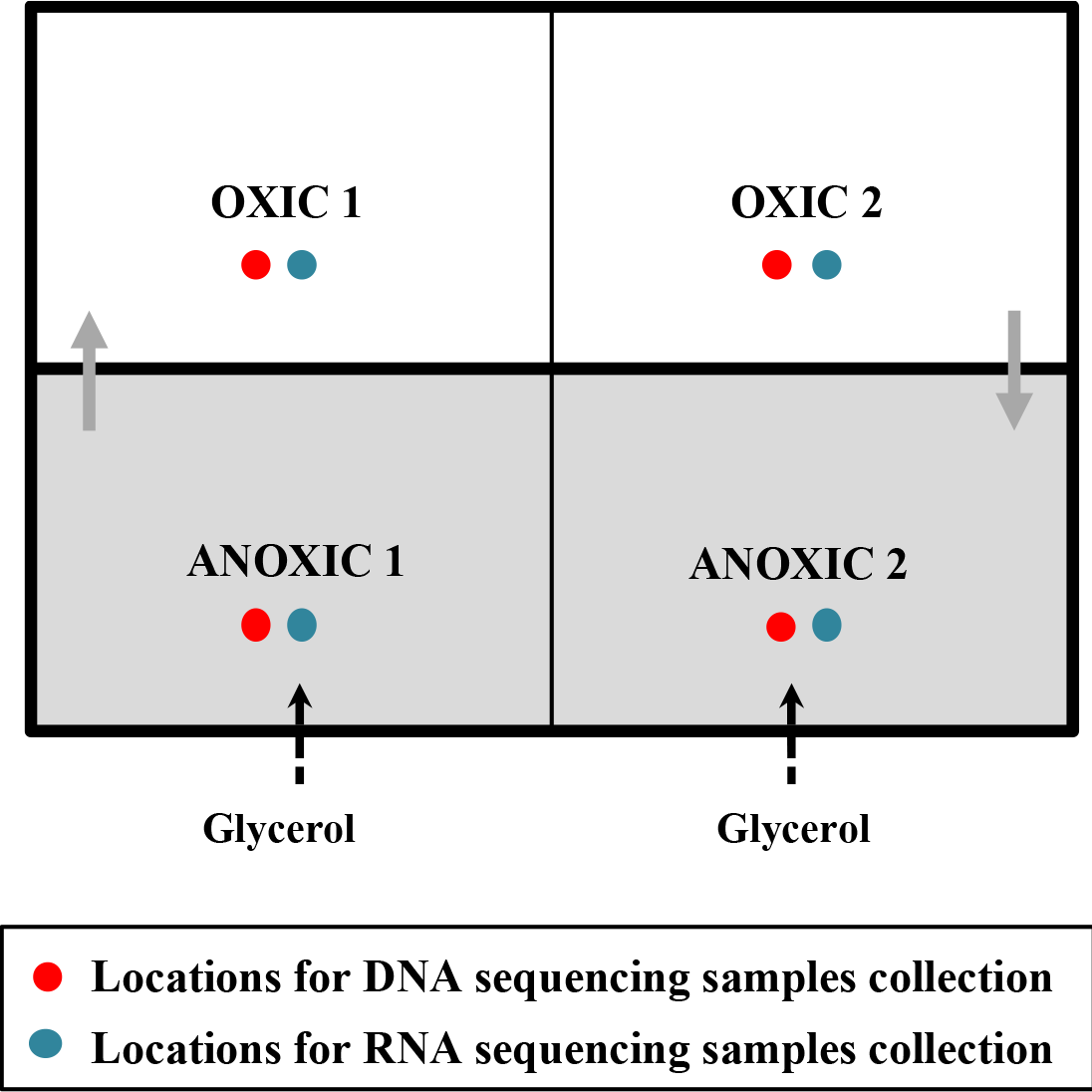
Schematic of SHARON process at Wards Island WPCP. Shaded areas represent anoxic zones. Green arrows indicate the flow of wastewater. Samples were collected for six consecutive months.

**Fig. S6.**
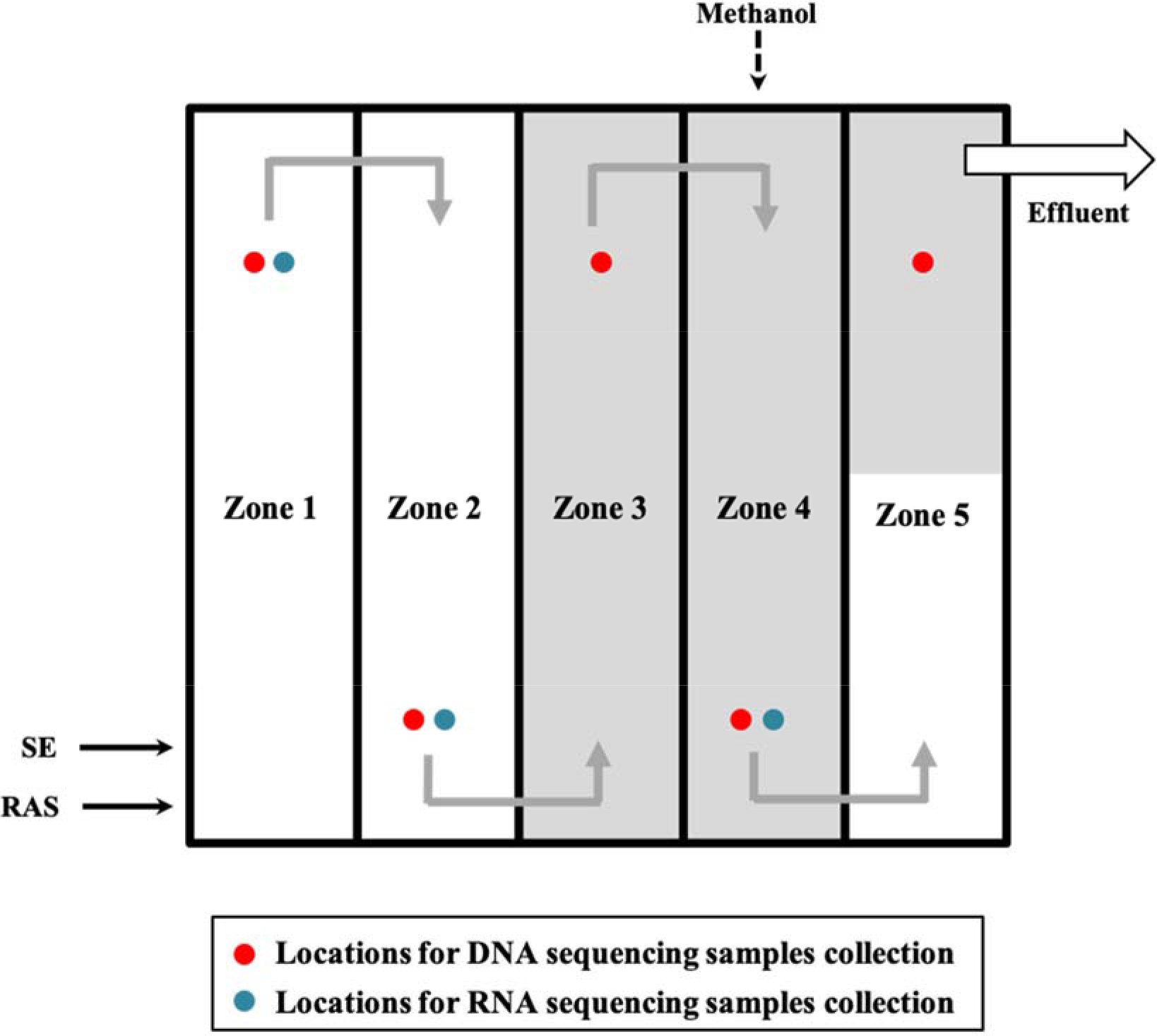
Schematic of DCW Blue Plains at advanced wastewater treatment plant. Shaded areas represent anoxic zones. SE: secondary effluent. Samples were collected for six consecutive months.

**Fig. S7.**
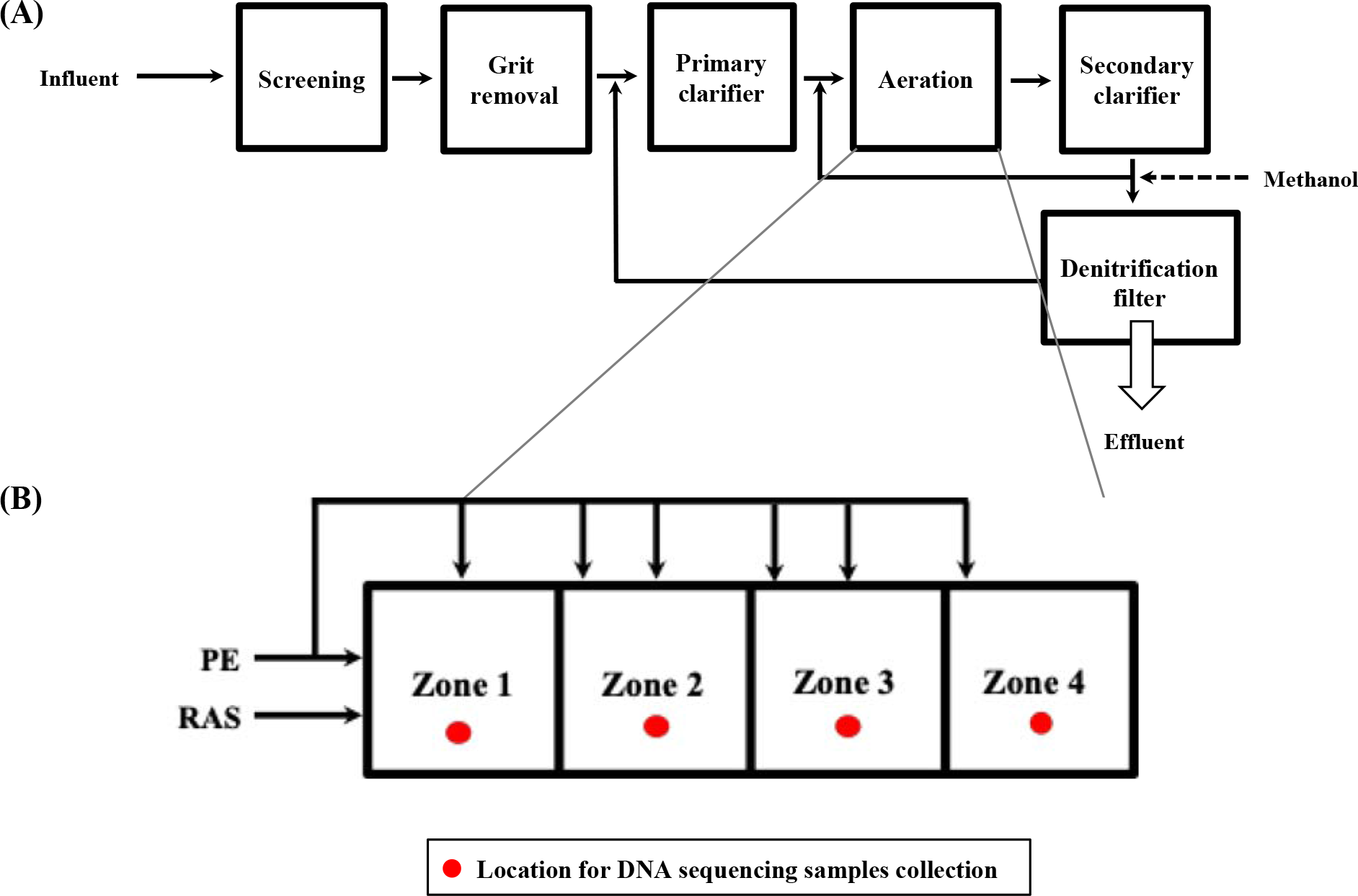
Schematic diagram of (A) York River treatment plant and (B) YRARE operating a step-feed aeration process. PE: primary effluent; RAS: return activated sludge. Samples were collected for six consecutive months.

**Fig. S8.**
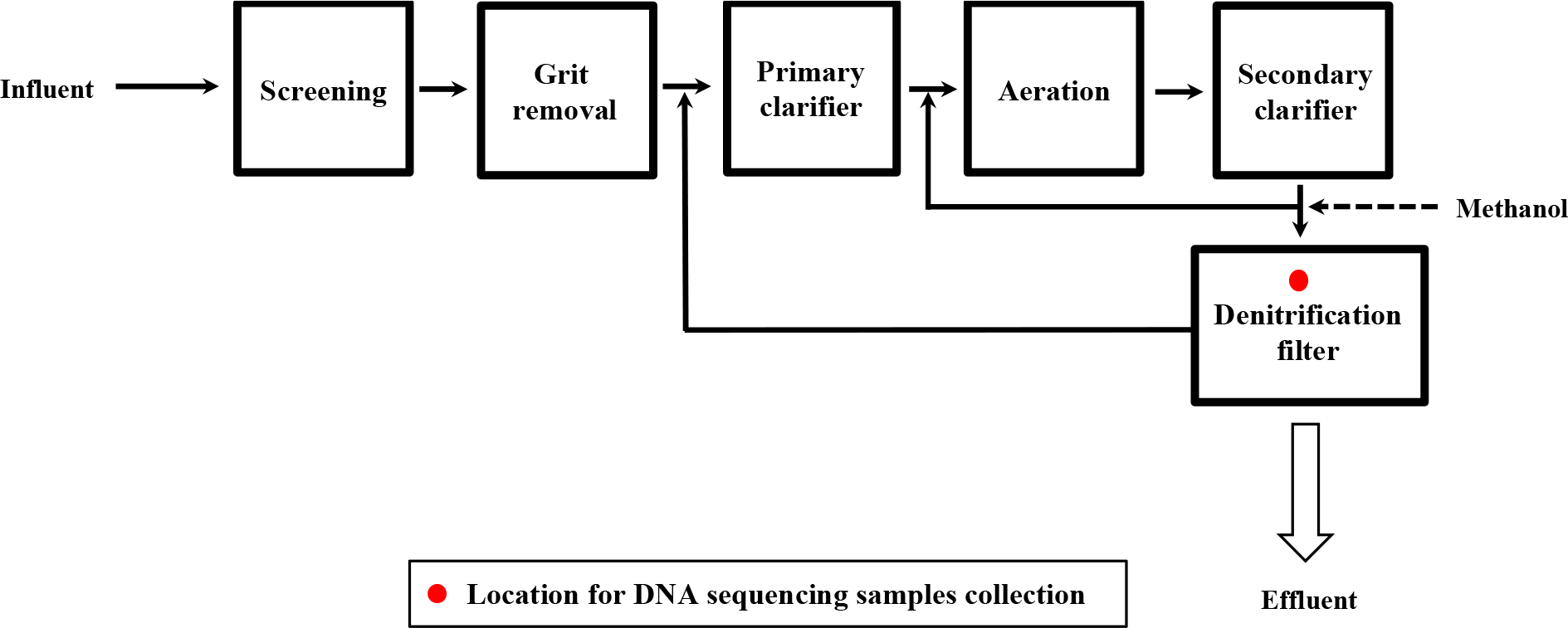
Schematic of YRDENI operating methanol-based denitrification. Samples were collected for six consecutive months.

**Fig. S9.**
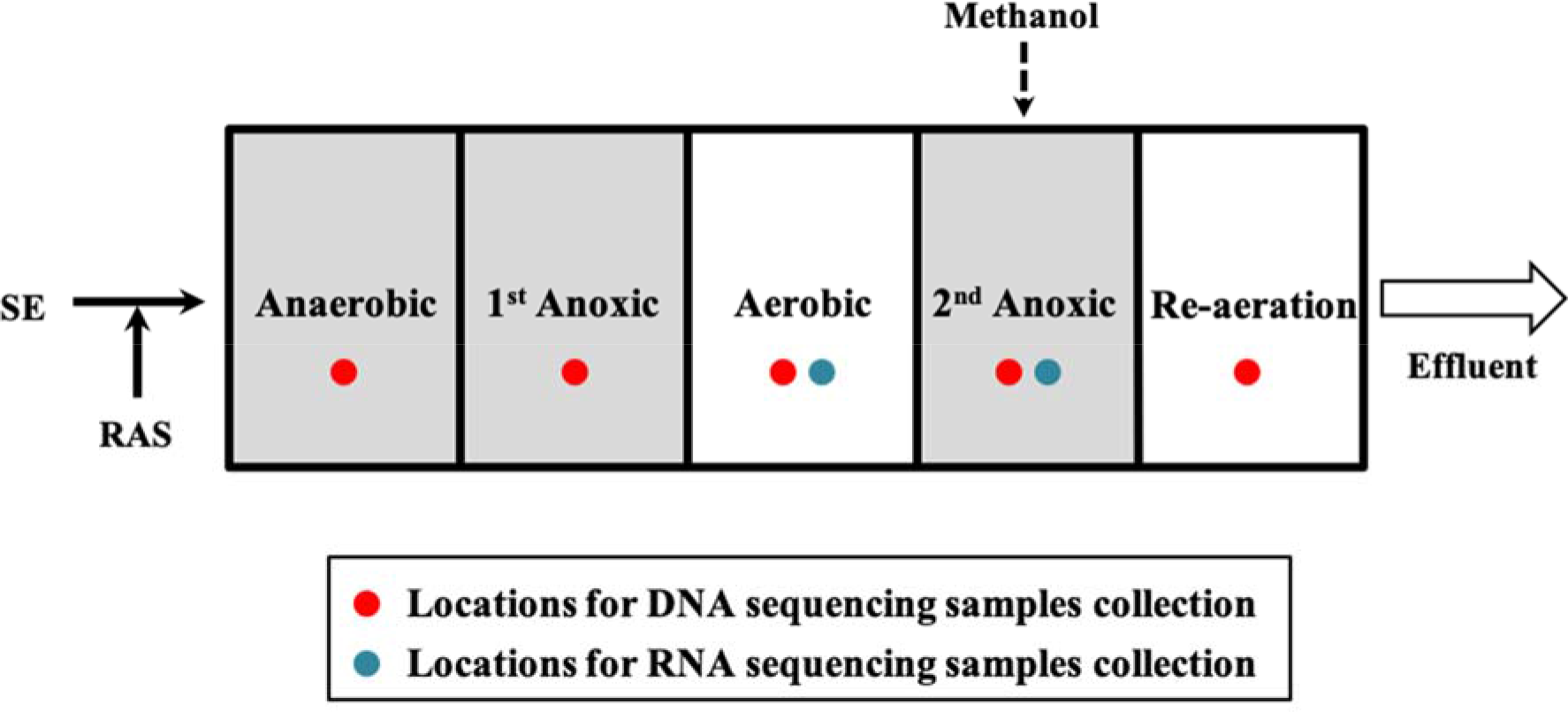
Schematic of NP operating 5-stage Bardenpho process. Shaded areas represent anoxic zones. Samples were collected for six consecutive months.

**Fig. S10.**
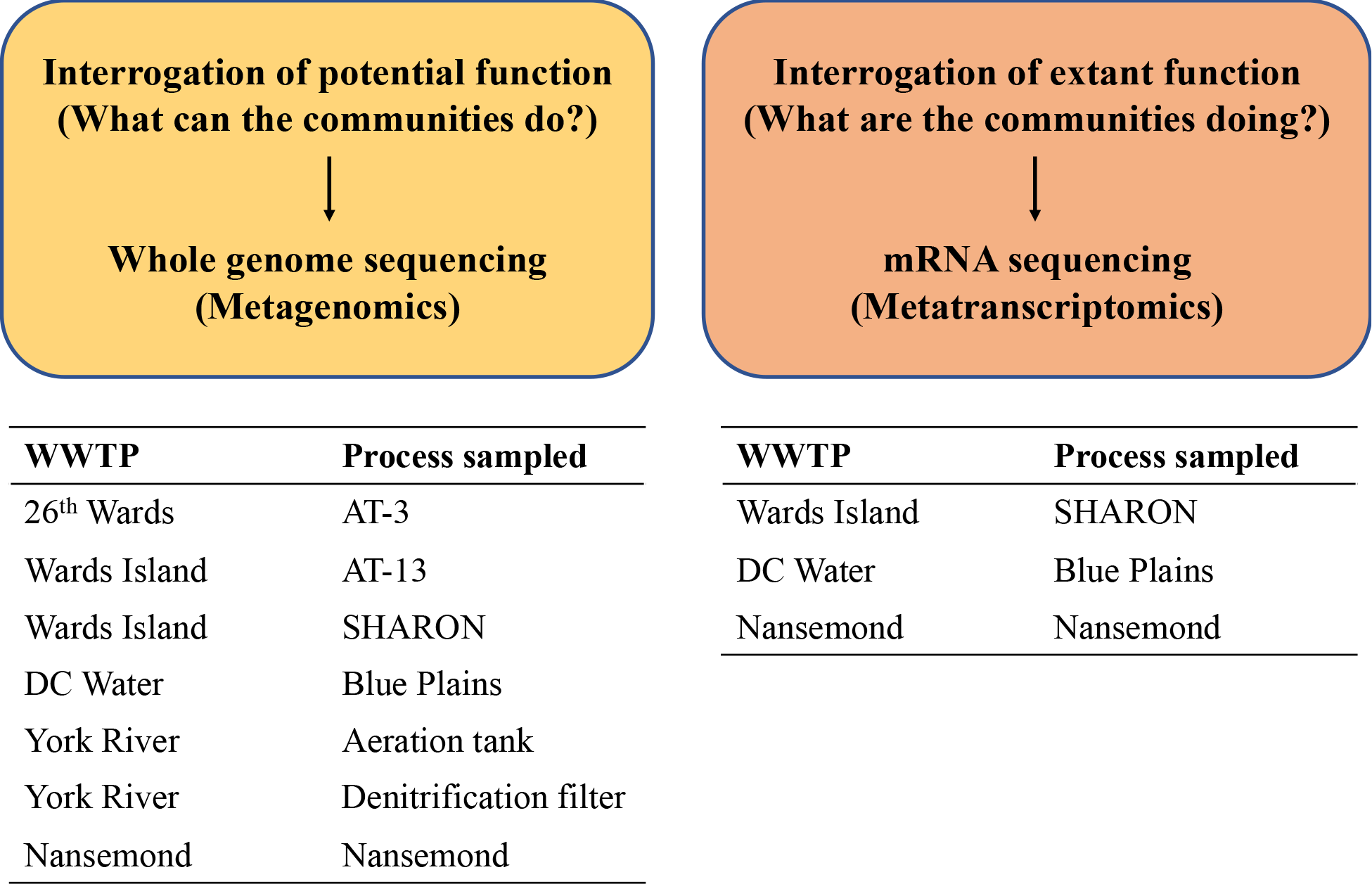
Schematic of meta-omics approach for interrogation of field-scale engineered wastewater treatment plants.

**Table S1.** Description of samples collected from seven process reactors at five WWTPs.

|  | **WWTP** | **Reactor** | **Configuration** | **Carbon source** | **Sampling period** |
| --- | --- | --- | --- | --- | --- |
| 1 | 26th Ward | AT-3 | Plug-flow separate centrate | Glycerol | Sep. 2014 - Mar. 2015<sup>a</sup> |
| 2 | Wards Island | AT-13 | Step-feed BNR | Glycerol |  |
| 3 |  | SHARON | Nitritation-denitritation | Glycerol |  |
| 4 | DC Water | Blue Plains | Post denitrification | Methanol | Nov. 2014 - Apr. 2015 |
| 5 | HRSD York River | York River (YR) | Nitrification | - | Nov. 2014 - Jun. 2015<sup>b</sup> |
| 6 |  |  | Denitrification filter | Methanol |  |
| 7 | HRSD Nansemond | Nansemond (NP) | 5-stage Bardenpho | Methanol |  |

<sup>a</sup>With the exception of Dec. 2014
<sup>b</sup>With the exception of Dec. 2014 and Feb. 2015

**Table S2.**
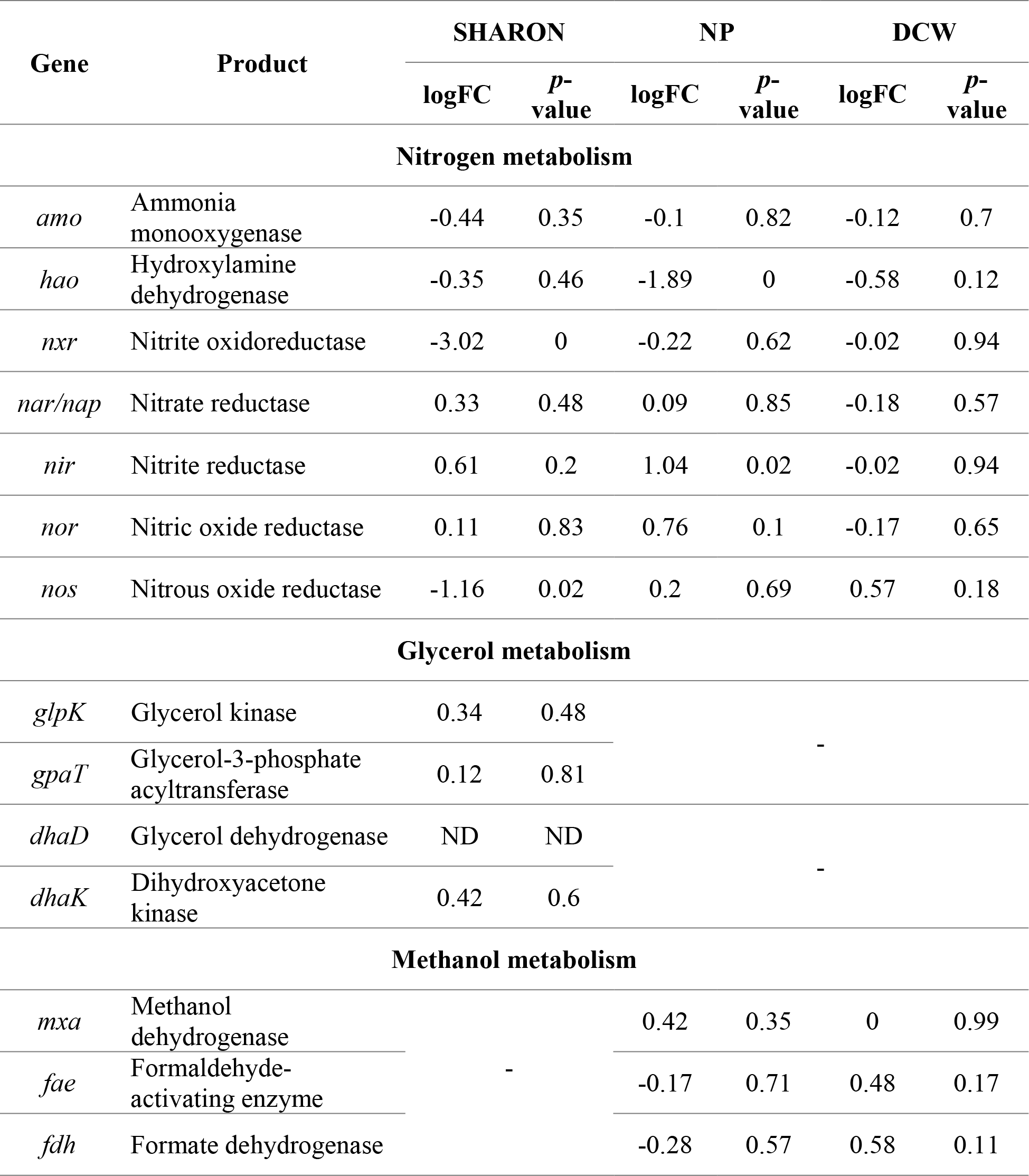
Log fold change (FC) and *p*-values of the differential gene expression analysis for the metatranscriptomes of SHARON, NP and DCW.

